# eDNA enables proof-of-origin of samples from reservoirs with rich and abundant biodiversity

**DOI:** 10.1101/2025.05.13.653644

**Authors:** Mauro Rebelo, Thainá Cortez, João Nunes Godinho, Juliana Coelho, Rosa Cavalcanti, Henrique Bomfim, Yasmin Cunha, Francesco Dondero, Danielle Amaral, Juliana Americo

## Abstract

Environmental DNA (eDNA) is increasingly used for biodiversity monitoring, but its application in verifying the geographic origin of samples remains limited. Here, we evaluated whether eDNA profiles from water samples can indicate sample provenance across three large Brazilian reservoirs. Using multi-marker metabarcoding (COI, 12S, 16S), we showed that biodiversity signatures enable spatial attribution. To ensure traceability, each report was encoded in a machine-readable format, hashed, and registered on the blockchain. This framework demonstrates a novel approach to independently verifying sample origin through molecular and cryptographic evidence.

## 1 Introduction

Over the past decade, eDNA metabarcoding has been increasingly adopted in ecological studies, conservation planning, and environmental licensing, particularly in aquatic ecosystems [1, 2].

Experimental applications of eDNA have demonstrated its potential to infer the geographic origin of environmental samples by analyzing the taxonomic composition of their microbial, fungal, or plant communities. In forensic investigations, metabarcoding of soil and dust has been used to attribute samples to their probable source regions with promising accuracy under controlled conditions [3] [4]. Similarly, in the field of public health, metagenomic profiling of microbial DNA from animal feces and water sources has been applied to assess cross-contamination and sanitation risks, linking microbial signatures to specific ecological and geographic contexts [5].

These studies underscore a growing interest in using eDNA as a proxy for biogeographic attribution. Yet, despite these advances, such applications remain largely experimental and are not yet integrated into standardized frameworks for conservation monitoring or regulatory enforcement. While the potential is clear, what remains lacking is a robust and auditable system that allows eDNA-based inferences to be used confidently in decision-making processes. Our work addresses this gap by proposing a decentralized and verifiable framework in which eDNA profiles serve as digital biomarkers of geographic origin.

We previously showed that saturation curves outperform environmental variables in validating sample completeness, both for individual (alpha) and grouped (beta) analyses [6].

Here, we replicate that approach in two other reservoirs that cover in total approximately 3,000 km2, using a standardized eDNA protocol with multiple genetic markers and sampling based on the reservoirs’ hydrodynamics that analyze Molecular Operational Taxonomic Units (MOTUs) comparison to taxonomic lists with BLAST to create biodiversity profiles.

We tested whether statistical analyses of richness and abundance, including discriminant and ordination methods, can assign each sample to its source reservoir based solely on its genetic profile. This would demonstrate that the reservoirs are sufficiently characterized for origin attribution and that eDNA can create a unique, reproducible biodiversity fingerprint.

If validated, this enables a new application for eDNA: sample provenance verification. Creates a trustless chain of custody that meets regulatory demands without relying on declarations, unlocking scalable, independent environmental monitoring.

## 2 Results and Discussion

### 2.1 Hydrodynamics explains eDNA patterns but does not constrain sampling design

Environmental DNA samples were collected from two hydroelectric reservoirs, Hydroelectric Power Plant (HPP) Jupiá and HPP Rosana, in southwestern Brazil (Figures 1a and 1b 1). Sampling campaigns were conducted in December 2022 (rainy season), November 2023 (transition to the rainy season), and July 2024 (the dry season). In the HPP Jupiá reservoir, 6 stations (JUP1–JUP6; Table 1) were sampled, while 9 stations (ROS1–ROS6, ROSE1–ROSE3) were also sampled in the HPP Rosana reservoir. The same stations were sampled in all campaigns, and additional control samples were collected to ensure data consistency. Station selection was guided by hydrodynamic modeling. The spatial distribution of sampling stations was mapped in both reservoirs. In HPP Jupiá, stations were distributed across upstream, central, and downstream regions, including major tributary in-flows. In HPP Rosana, stations were positioned to cover a range of hydrologic conditions, from high-flow inflows and outflows to low-velocity mid-reservoir zones.

**Table 1.** Geolocation and sampling dates of the environmental DNA (eDNA) collection sites in the Rosana and Jupiá reservoirs. The table lists the latitude and longitude of each sampling station, along with the corresponding collection dates for each survey event.

| **Sampling Station** | **Latitude** | **Longitude** | **Sampling Date** |
| --- | --- | --- | --- |
| **Survey 1 (December 2022)** |  |  |  |
| ROS#01 | -22.598959 | -52.848337 | 2022-12-13 |
| JUP#01 | -20.761749 | -51.633791 | 2022-12-14 |
| **Survey 2 (November 2023)** |  |  |  |
| ROS#01 | -22.606409 | -52.861704 | 2023-11-16 |
| ROS#02 | -22.575588 | -52.591398 | 2023-11-01 |
| ROS#03 | -22.628626 | -52.498430 | 2023-11-16 |
| ROS#04 | -22.671562 | -52.227803 | 2023-11-17 |
| ROS#05 | -22.610286 | -52.159356 | 2023-11-17 |
| ROS#06 | -22.557507 | -52.149941 | 2023-11-17 |
| JUP#03 | -20.639504 | -51.589322 | 2023-11-13 |
| JUP#02 | -20.680354 | -51.712669 | 2023-11-13 |
| JUP#01 | -20.759253 | -51.630821 | 2023-11-13 |
| JUP#04 | -20.666945 | -51.484590 | 2023-11-13 |
| JUP#05 | -20.525337 | -51.480978 | 2023-11-14 |
| JUP#06 | -20.407516 | -51.380272 | 2023-11-14 |
| **Survey 3 (July 2024)** |  |  |  |
| ROS#01 | -22.60000 | -52.84806 | 2024-07-29 |
| ROS#02 | -22.58333 | -52.58611 | 2024-07-29 |
| ROS#03 | -22.63167 | -52.48556 | 2024-07-29 |
| ROS#04 | -22.67778 | -52.22028 | 2024-07-30 |
| ROS#05 | -22.61389 | -52.16000 | 2024-07-30 |
| ROS#06 | -22.56111 | -52.15028 | 2024-07-30 |
| ROSE#01 | -22.62500 | -52.69694 | 2024-07-29 |
| ROSE#02 | -22.61500 | -52.54778 | 2024-07-29 |
| ROSE#03 | -22.61778 | -52.24583 | 2024-07-29 |
| JUP#01 | -20.76389 | -51.63000 | 2024-07-25 |
| JUP#02 | -20.69444 | -51.70083 | 2024-07-25 |
| JUP#03 | -20.63750 | -51.58611 | 2024-07-25 |
| JUP#04 | -20.66444 | -51.50361 | 2024-07-25 |
| JUP#05 | -20.51667 | -51.46722 | 2024-07-26 |
| JUP#06 | -20.41667 | -51.37500 | 2024-07-26 |

**Figure 1.**
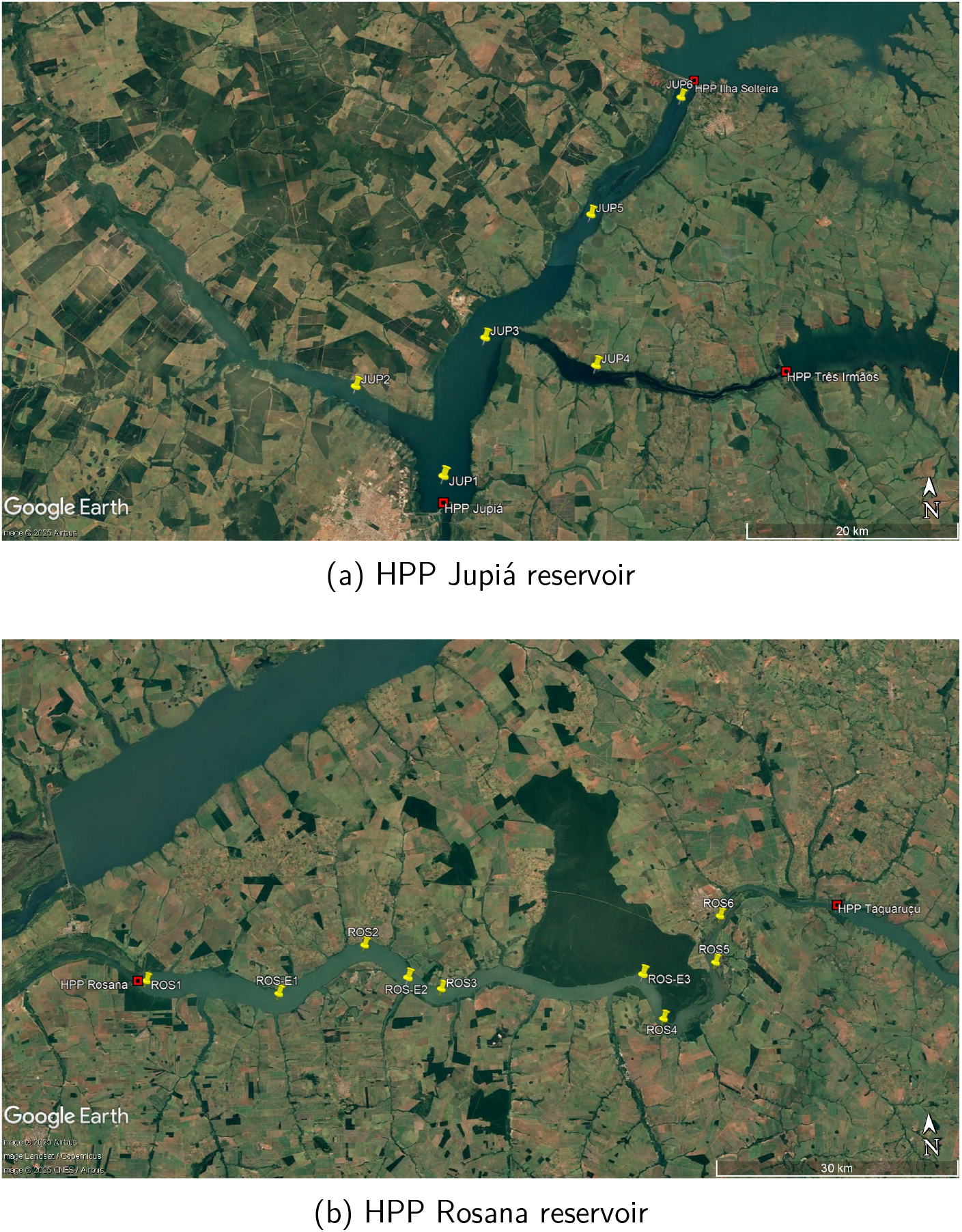
Satellite image showing the spatial distribution of environmental DNA sampling stations in the (a) UHE Jupiá reservoir and (b) UHE Rosana reservoir. Points span up, mid, and downstream locations. Geolocation coordinates are provided in Table 1. Projection: UTM. Datum: SIRGAS2000.

The hydrodynamic simulation results for the Jupiá and Rosana Reservoirs revealed distinct flow regimes that have implications for the distribution and retention of environmental DNA (eDNA).

In the Jupiá Reservoir, during the rainy season, time series data recorded maximum flow velocities of 0.10 m/s at the upstream inflows from the Ilha Solteira and Três Irmãos HPPs, while the central reservoir maintained velocities below 0.02 m/s. On simulation day 87, the instantaneous current field reached a peak of 0.128 m/s along the main channel of the Paraná River. In the dry season, maximum velocities near the upstream inflows were 0.07 m/s, with other stations recording values between 0.015 and 0.035 m/s, and an instantaneous maximum of 0.122 m/s was observed on day 85 along the upstream channel. Further analysis on day 87 during the rainy season indicated that within the Paraná River channel, velocities ranged from 0.024 to 0.067 m/s, with a mean of 0.064 m/s (± 0.014) immediately downstream of the Ilha Solteira HPP; in the Tietê River, a maximum of 0.05 m/s was observed; in the Sucuriú River, the maximum velocity was 0.021 m/s; and near the Jupiá HPP in the downstream area, velocities reached 0.076 m/s. These measurements, which are relatively low compared to typical riverine flows exceeding 1 m/s, indicate that the Jupiá Reservoir exhibits a predominantly lentic environment in its central regions, with moderately faster flows confined to the inflow and main channel areas.

In contrast, the Rosana Reservoir exhibited considerably higher velocities. During the dry season, time series data showed an upstream peak of 3.1 m/s on day 38, with the instantaneous current field indicating strong upstream flow while central and western areas maintained velocities below 1.0 m/s. In the rainy season, time series data recorded upstream velocities of up to 2.4 m/s and downstream velocities of 1.6 m/s, and on day 50, the instantaneous current field reached 2.5 m/s along the eastern main channel. Further spatial analysis on the rainy season day 50 revealed that near HPP Taquaruçu, upstream velocities reached 2.4 m/s with an average of 1.15 m/s, whereas near HPP Rosana in the downstream area, velocities exceeded 1 m/s, and a cyclonic recirculation pattern was observed. The central channel exhibited low velocities with a modal value of approximately 0.1 m/s, and the meandering channel showed intermediate velocities ranging from 0.45 to 0.74 m/s, likely as a result of channel narrowing and curvature.

The contrasting hydrodynamic conditions between the two reservoirs directly influence eDNA dynamics. In the Jupiá Reservoir, the low velocities in the central, lentic regions promote local retention and accumulation of eDNA due to extended residence times. However, in the moderately faster inflow and channel zones, advection drives downstream transport of both dissolved and particle-attached DNA, potentially integrating signals from upstream sources. In the Rosana Reservoir, the substantially higher velocities in the upstream regions and main channels facilitate rapid advective transport of eDNA over longer distances. Nonetheless, areas with low flow, such as the central channel and zones exhibiting cyclonic recirculation, may foster local retention and sedimentation of particle-attached DNA. These spatial variations imply that eDNA sampled in high-velocity zones is likely to reflect contributions from distant upstream sources, whereas samples from low-velocity areas yield a more localized signal.

Integrating the hydrodynamic and eDNA data from both reservoirs underscores the importance of considering spatial heterogeneity and seasonal variability when designing eDNA monitoring strategies. In systems like Jupiá, with predominantly low velocities, local biological communities are more directly represented in eDNA samples, while in Rosana, the dynamic range of flow conditions requires careful spatial sampling to distinguish between locally produced and advected eDNA. This integrated approach supports the interpretation of eDNA signals in relation to the physical transport processes, thereby enhancing the accuracy of biodiversity assessments, invasive species detection, and studies on organism dispersal in reservoir environments.

The integrated analysis suggests that while hydrodynamic modeling provides detailed insight into eDNA transport processes, its cost and labor intensity may not justify its routine application for broad biodiversity assessments. The biodiversity results indicate that random sampling alone can effectively capture the overall species distribution across both reservoirs. Thus, for many monitoring programs, a cost-effective approach may be to rely primarily on random sampling, supplemented by targeted hydrodynamic analysis only when fine-scale spatial resolution of eDNA sources is essential.

### 2.2 Bioinformatics shows the quality of the DNA sampling, extraction and sequencing

The sequencing data underwent standard quality control and processing steps, including filtering, denoising, merging, and chimera removal.

The analysis revealed varying sequencing depths and quality metrics across campaigns and reservoirs**??**. Total reads ranged from approximately 133,000 to over 2.8 million per reservoir per campaign. For the third survey in July 2024, we successfully identified multiple genetic markers (COI, 12S, and 16S), resulting in 2,530 ASVs and 1,804 OTUs for COI, 449 ASVs and 357 OTUs for 12S, and 778 ASVs and 416 OTUs for 16S.

Although OTU clustering was performed only for the third survey, the richness of ASVs and OTUs detected (Table 3) highlights the effectiveness and robustness of our multi-marker molecular approach in capturing biodiversity.

**Table 2.** The table presents key sequencing metrics, including the total number of reads, reads retained after quality filtering, percentage of reads passing the filter, denoised reads, merged reads, and non-chimeric sequences. Data are shown for three sampling campaigns conducted in December 2022 (1^st^), November 2023 (2^nd^), and July 2024 (3^rd^). The percentage of non-chimeric sequences reflects the proportion of high-quality reads available for taxonomic classification.

| Reservoir | Survey | Total Reads | Filtered | % Passed | Denoised | Merged | Non-Chim. | % Non-Chim. |
| --- | --- | --- | --- | --- | --- | --- | --- | --- |
| Jupiá | 1st | 2,887,380 | 2,816,406 | 97.54 | 2,810,859 | 615,942 | 608,723 | 21.73 |
| Jupiá | 2nd | 392,521 | 377,323 | 96.13 | 376,189 | 27,683 | 27,538 | 6.73 |
| Jupiá | 3rd | 426,095 | 357,894 | 83.99 | 357,081 | 10,270 | 10,270 | 2.41 |
| Rosana | 1st | 2,367,431 | 2,305,698 | 97.39 | 2,295,212 | 1,268,761 | 1,259,121 | 53.19 |
| Rosana | 2nd | 133,110 | 126,504 | 95.04 | 126,101 | 87,292 | 86,337 | 64.86 |
| Rosana | 3rd | 214,649 | 181,265 | 84.45 | 180,852 | 4,486 | 4,486 | 2.45 |

**Table 3.** Number of Amplicon Sequence Variants (ASVs) and Operational Taxonomic Units (OTUs) detected for each genetic marker during the third sampling campaign in the Jupiá and Rosana reservoirs.

| Survey | Reservoir | Genetic Marker | ASVs | OTUs |
| --- | --- | --- | --- | --- |
| 3rd | Jupiá | COI | 2530 | 1804 |
| 3rd | Jupiá | 12S | 449 | 357 |
| 3rd | Jupiá | 16S | 778 | 416 |
| 3rd | Rosana | COI | 1904 | 1395 |
| 3rd | Rosana | 12S | 888 | 434 |
| 3rd | Rosana | 16S | 857 | 492 |

Further details on the taxonomic classification obtained through our multimarker molecular approach during the third campaign (Rosana and Jupiá), including comprehensive classification tables for each genetic marker used, are provided in supplementary material 1, 2 e 3

**Figure 2.**
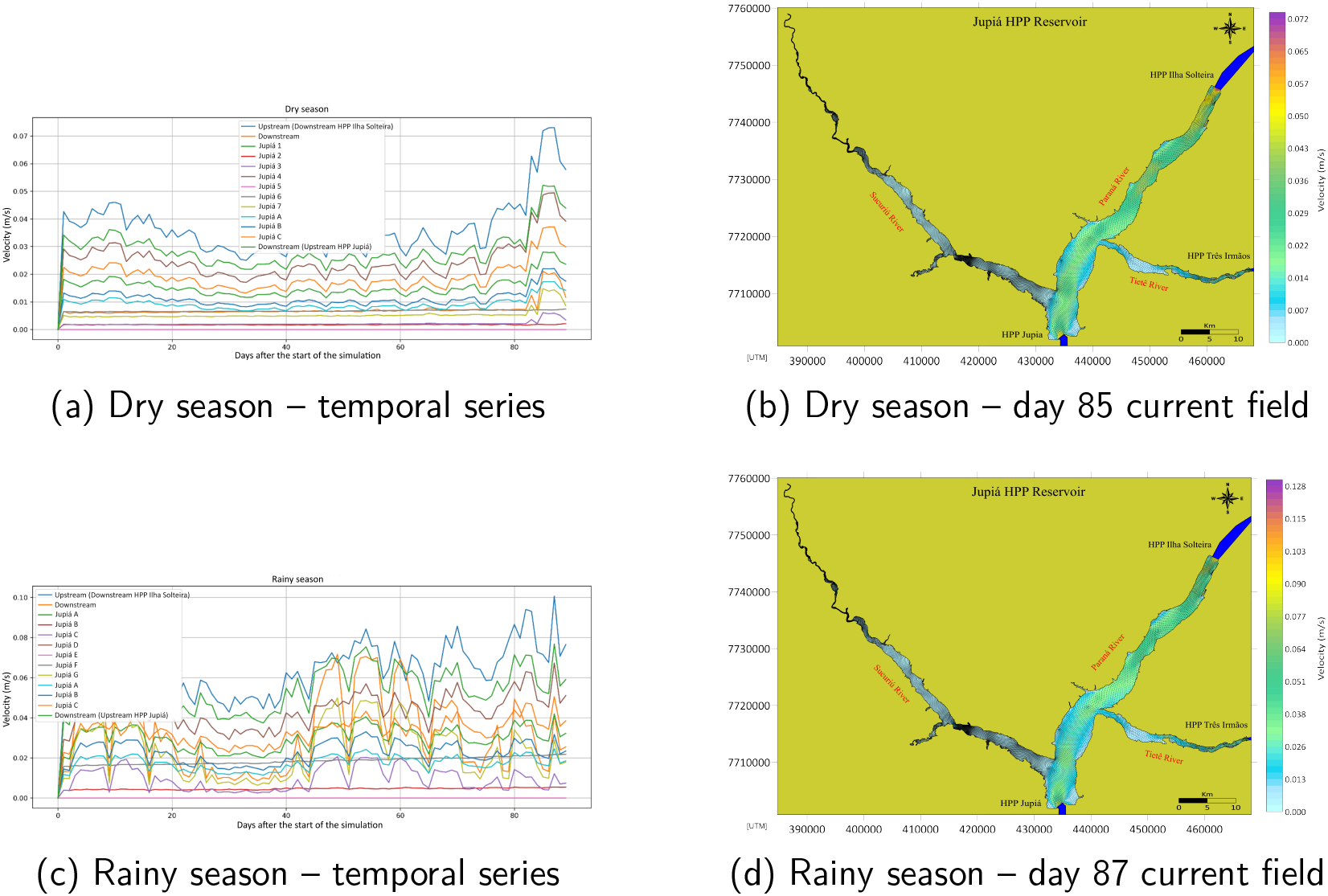
Hydrodynamics in the Jupiá Reservoir. (a) Time series of flow velocity at key sampling points during the dry season, showing lower and more stable velocities. (b) Instantaneous current field on day 85 (dry season maximum), with restricted fast flow near the inflows and stagnation elsewhere. (c) Time series of flow velocity during the rainy season, with peaks near upstream inflows ( 0.10 m/s). (d) Instantaneous current field on day 87 (rainy season maximum), showing peak values (up to 0.128 m/s) in the main channel. Projection: UTM. Datum: SIRGAS2000.

**Figure 3.**
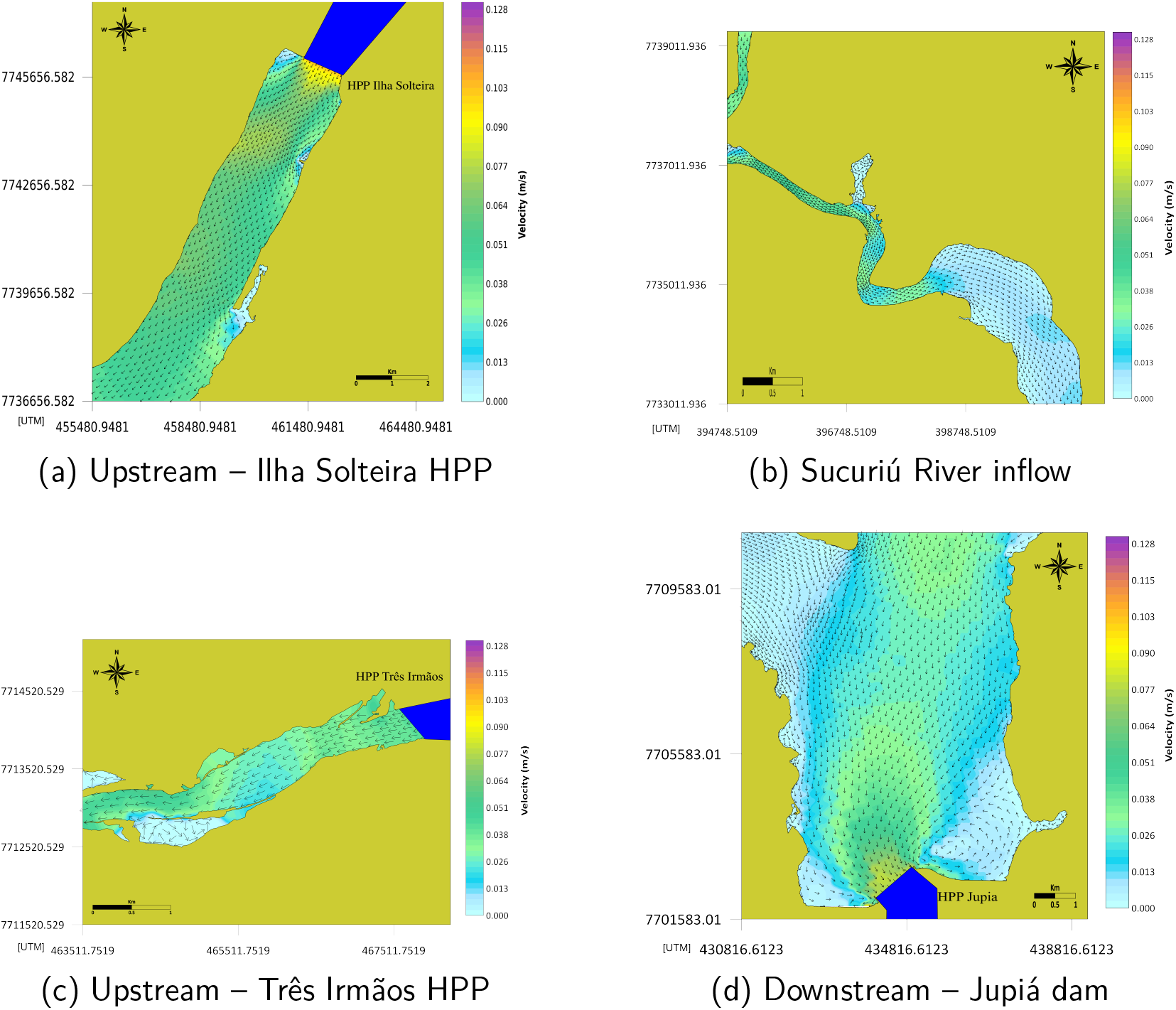
Velocity and current fields in the Jupiá Reservoir on day 87 (rainy season). (a) Upstream region near Ilha Solteira HPP: max 0.1 m/s, avg 0.064 ± 0.014 m/s. (b) Sucuriú River inflow: extremely low velocities (¡0.021 m/s). (c) Upstream near Três Irmãos HPP: flow ¡0.05 m/s. (d) Downstream region near Jupiá dam: local peaks up to 0.076 m/s. Projection: UTM. Datum: SIRGAS2000.

**Figure 4.**
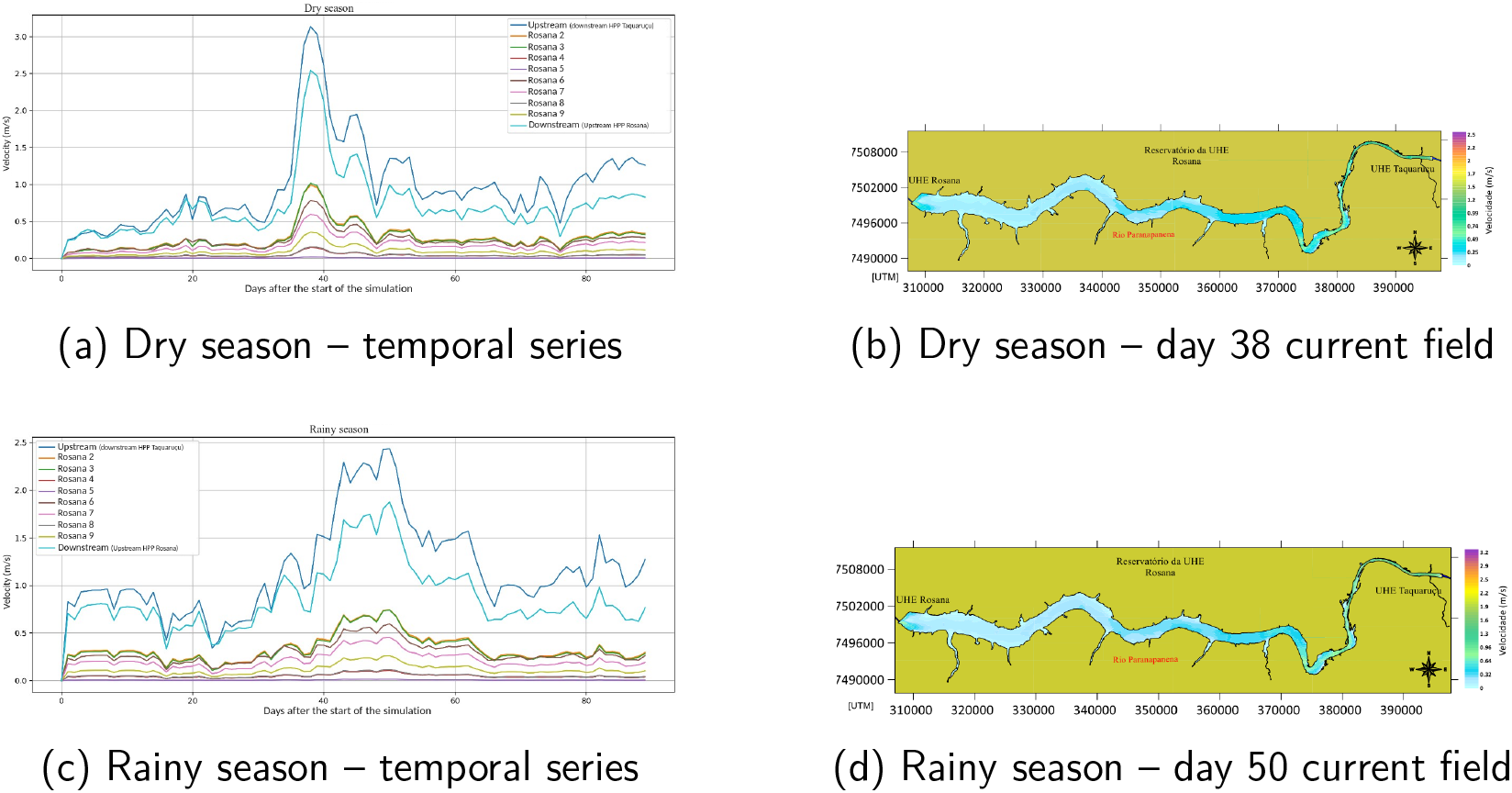
Hydrodynamics in the Rosana Reservoir. (a) Time series of flow velocity during the dry season, with upstream peak at 3.1 m/s on day 38. (b) Instantaneous current field on day 38 (dry season maximum), showing strong upstream flow and slower central/western areas (¡1.0 m/s). (c) Time series of flow velocity during the rainy season, showing peaks up to 2.4 m/s upstream and 1.6 m/s downstream. (d) Instantaneous current field on day 50 (rainy season maximum), with flow reaching 2.5 m/s along the eastern main channel. Projection: UTM. Datum: SIRGAS2000.

**Figure 5.**
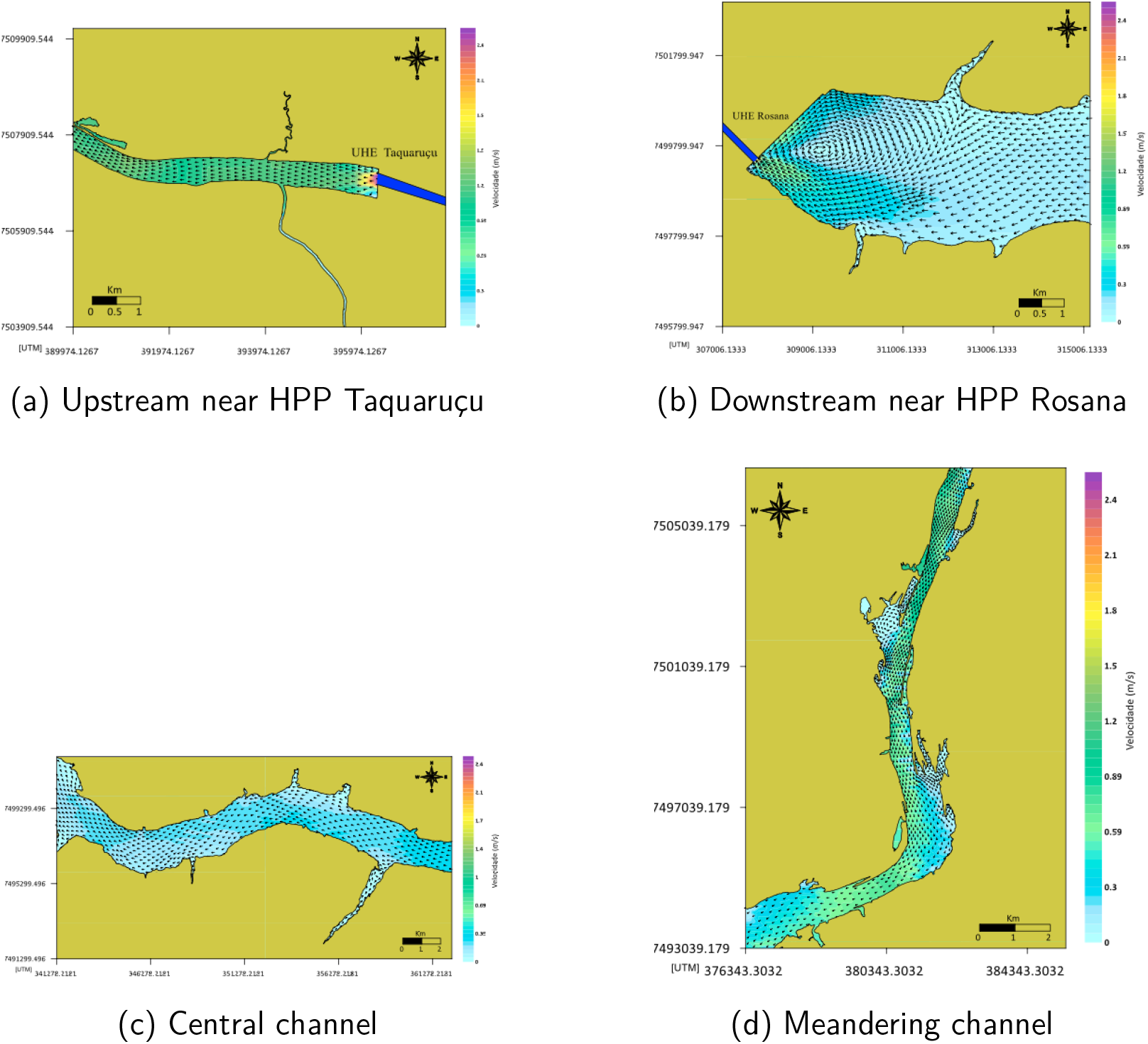
Velocity and current fields in the Rosana Reservoir (rainy season, day 50). (a) Upstream near HPP Taquaruçu: velocities up to 2.4 m/s, average 1.15 m/s. (b) Downstream near HPP Rosana: velocities ¿1 m/s and visible cyclonic recirculation. (c) Central channel: low velocity zone with modal value 0.1 m/s. (d) Meandering channel: intermediate velocities (0.45–0.74 m/s) due to narrowing and curvature. Projection: UTM. Datum: SIRGAS2000.

### 2.3 Taxonomic Composition and Conservation Status of Fish Communities in Jupiá Reservoir

A total of 220 amplicon sequence variants (ASVs) were identified across the three sampling campaigns in the Jupiá reservoir, representing 10 phyla, 26 classes, 57 orders, 96 families, 90 genera, and 138 species (figure 6). The complete list is provided in the supplementary materials 4. Eigth species, 16 genera, and 14 families were consistently detected in all three surveys.

**Figure 6.**
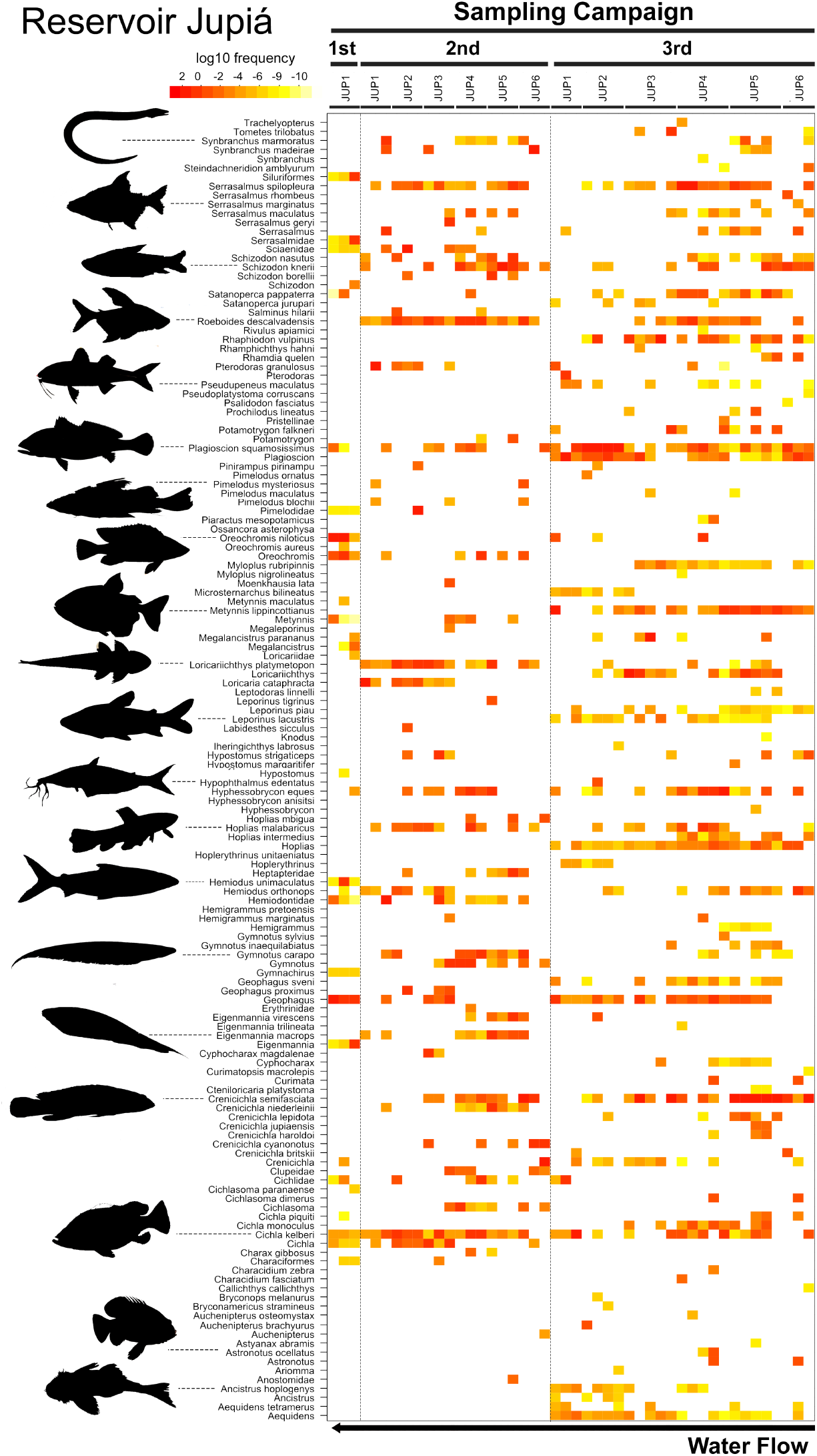
Heatmap of fish species diversity and abundance in the Jupiá Reservoir, for all 3 primers, cumulative in all 3 surveys

These numbers may reflect identification biases inherent to molecular methods in which species-level assignment depends on the availability of reference sequences in public databases. In cases where species-level identification was not possible, sequences were assigned to the closest available taxonomic rank, which may result in a higher number of genera than families.

The detection of **16 genera** across all sampling campaigns suggests the existence of a core group of taxa that persist in the reservoir, despite environmental variability or sampling differences. The lower number of consistently detected species (8) may reflect limitations in marker resolution, DNA degradation, or sequencing depth.

We identified **10 species** as native to the Paraná River Basin, expected components of the regional ichthyofauna. These include: *Astyanax lacustris* (lambari), *Leporinus obtusidens* (piapara), *Brycon orbignyanus* (piracanjuba), *Salminus brasiliensis* (dourado) *Pimelodus maculatus* (mandi), *Pseudoplatystoma corruscans* (surubim, pintado), *Prochilodus lineatus* (curimbatá), *Rhamdia quelen* (jundiá)

Four species exhibited broad geographic distribution and are considered cosmopolitan in South America: *Hoplias malabaricus* (traíra), *Ctenobrycon spilurus* (tetra prateado), *Hemigrammus marginatus*(piaba), and *Gymnotus carapo* (sarapÓ). Their presence is consistent with their known range across the Paraná, Amazon, São Francisco, Tocantins-Araguaia, and Orinoco basins.

Nine species were considered false positives, likely due to misclassification stemming from database limitations or sequence similarity with taxa from biogeographically distant systems.

We also detected seven exotic or invasive species, known to be introduced by human activities, particularly aquaculture and sport fishing: *Cichla ocellaris* (tucunaré-amarelo), *Cichla monoculus* (tucunaré), *Plagioscion squamosissimus* (pescada-do-Piaúi), *Colossoma macropomum* (tambaqui), *Oreochromis niloticus* (tilápia-do-Nilo), *Coptodon rendalli* (tilápia-vermelha), *Cyprinus carpio* (carpa).

These introductions raise concern due to their potential to impact native fish populations through predation, competition, habitat modification, and hybridization. For example, *Cichla* spp. are top predators, and tilápia species are known to degrade habitats and displace native cichlids.

Regarding conservation status, no species identified in this study were classified as Critically Endangered (CR) according to the IUCN Red List [7]. However, one species, *Steindachneridion melanodermatum* (jaú or surubim-do-Iguaçu), is categorized as Endangered (EN), primarily due to overfishing. Its detection in the reservoir is encouraging, suggesting that some populations persist.

Two species were listed as Vulnerable (VU): *Brycon orbignyanus* (piracanjuba) — migratory, impacted by fishing and damming, and *Pseudoplatystoma corruscans* (pintado or surubim-capivara), commercially valuable, declining.

All remaining species fell under Least Concern (LC) or Near Threatened (NT), indicating they are not currently at high extinction risk but may experience local pressures. Continued monitoring is essential to ensure sustainable management and conservation of fish biodiversity in the Paraná River basin.

### 2.4 Taxonomic Composition and Conservation Status of Fish Communities in Rosana Reservoir

In the Rosana reservoir, the environmental DNA (eDNA) survey detected a total of 217 amplicon sequence variants (ASVs) across all three sampling campaigns. These represented a broad taxonomic diversity, comprising 11 phyla, 24 classes, 52 orders, 93 families, 88 genera, and 133 species. The complete taxonomic classification is available in the supplementary material 5.

A consistent subset of 10 species, 15 genera, and 13 families was identified across all sampling campaigns, indicating the presence of a core ichthyofaunal community within the reservoir. This core likely reflects stable or resident species that persist across seasons and hydrological changes. As observed for Jupiá, discrepancies across taxonomic levels (e.g., more genera than families) are attributable to gaps in reference databases and limitations in species-level resolution of molecular markers.

The native fish community included multiple species endemic or typical of the Paraná River Basin. Notably, *Astyanax lacustris, Prochilodus lineatus* (curimbatá), *Pimelodus maculatus* (mandi), and *Leporinus obtusidens* (piapara) were consistently detected and are considered important ecological components of the basin.

Several cosmopolitan species, such as *Hoplias malabaricus* (traíra), *Gymnotus carapo* (saparÓ), and *Ctenobrycon spilurus* (tetra prateado), were also identified. These taxa are common across various South American basins and are not indicative of recent introductions or invasions.

As in Jupiá, the analysis revealed the presence of invasive and exotic species. Among the most notable were: *Cichla ocellaris* and *Cichla monoculus* (tucunarés) — introduced from the Amazon for sport fishing; *Oreochromis niloticus* and *Coptodon rendalli* (tilápias) — widespread aquaculture introductions and *Cyprinus carpio* (carpa) and *Colossoma macropomum* (tambaqui). These species are known to alter food webs and disrupt native communities through competition, predation, and habitat modification. Their detection supports the use of eDNA for early identification and monitoring of biological invasions.

In terms of conservation status, one Endangered (EN) species, *Steindachneridion melanodermatum* (surubim-do-iguaçu), was identified. Its detection in both reservoirs reinforces the potential of eDNA to support conservation by confirming the persistence of Endangered or Vulnerable taxa. Additionally, two species were listed as Vulnerable (VU) by the IUCN [7]: *Brycon orbignyanus* (piracanjuba) and *Pseudoplatystoma corruscans* (pintado). All remaining species were classified as Least Concern (LC) or Near Threatened (NT), with no species falling under the Critically Endangered (CR) category. This pattern mirrors that observed in Jupiá and reflects both regional conservation patterns and the effectiveness of eDNA for broad-scale biodiversity assessment.

**Figure 7.**
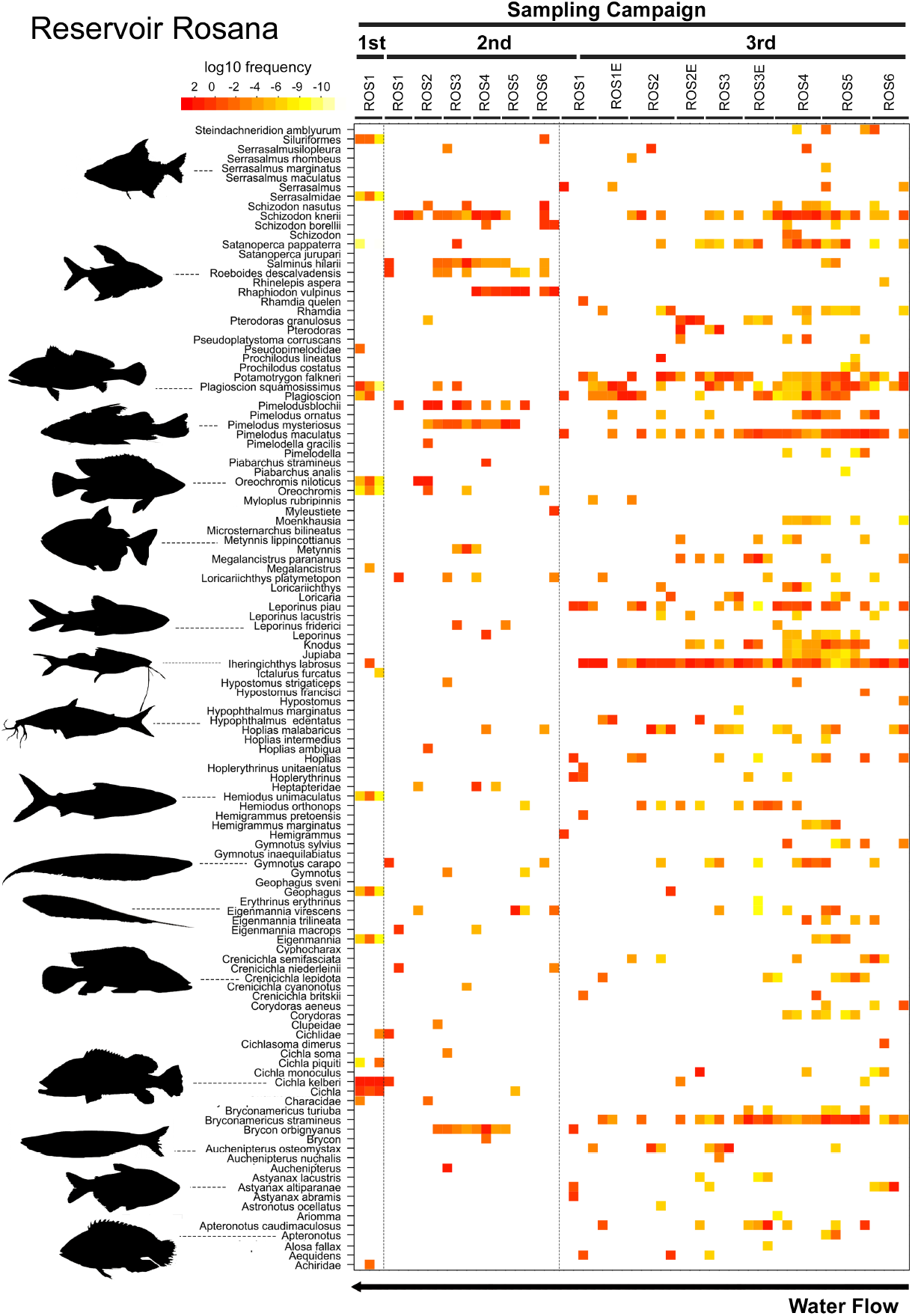
Heatmap of fish species diversity and abundance in the Rosana Reservoir, for all 3 primers, cumulative in all 3 surveys

### 2.5 COI is an effective marker for resolving genetic differences among the reservoirs

To estimate the ability of molecular biodiversity profiles to attribute provenance, we included data previously published from the Três Irmãos reservoir ([6]) to compare with current data from Jupiá and Rosana Reservoirs. Among the three genetic markers used in this study (COI, 12S, and 16S), the cytochrome oxidase I (COI) gene proved to be the most informative for distinguishing biodiversity patterns between the Rosana, Jupiá, and Três Irmãos reservoirs.

The COI marker consistently generated higher richness metrics across campaigns. In the third campaign (July 2024), it yielded 2,530 ASVs and 1,804 OTUs, compared to 449 ASVs / 357 OTUs for 12S and 778 ASVs / 416 OTUs for 16S in Jupiá and 1,904 ASVs and 1,395 OTUs, compared to 888 ASVs / 434 OTUs for 12S and 857 ASVs / 492 OTUs for 16S in Rosana(see Table 3). This higher resolution was evident in both taxonomic classification and the capacity to separate samples by reservoir of origin in multivariate analyses.

**Figure 8.**
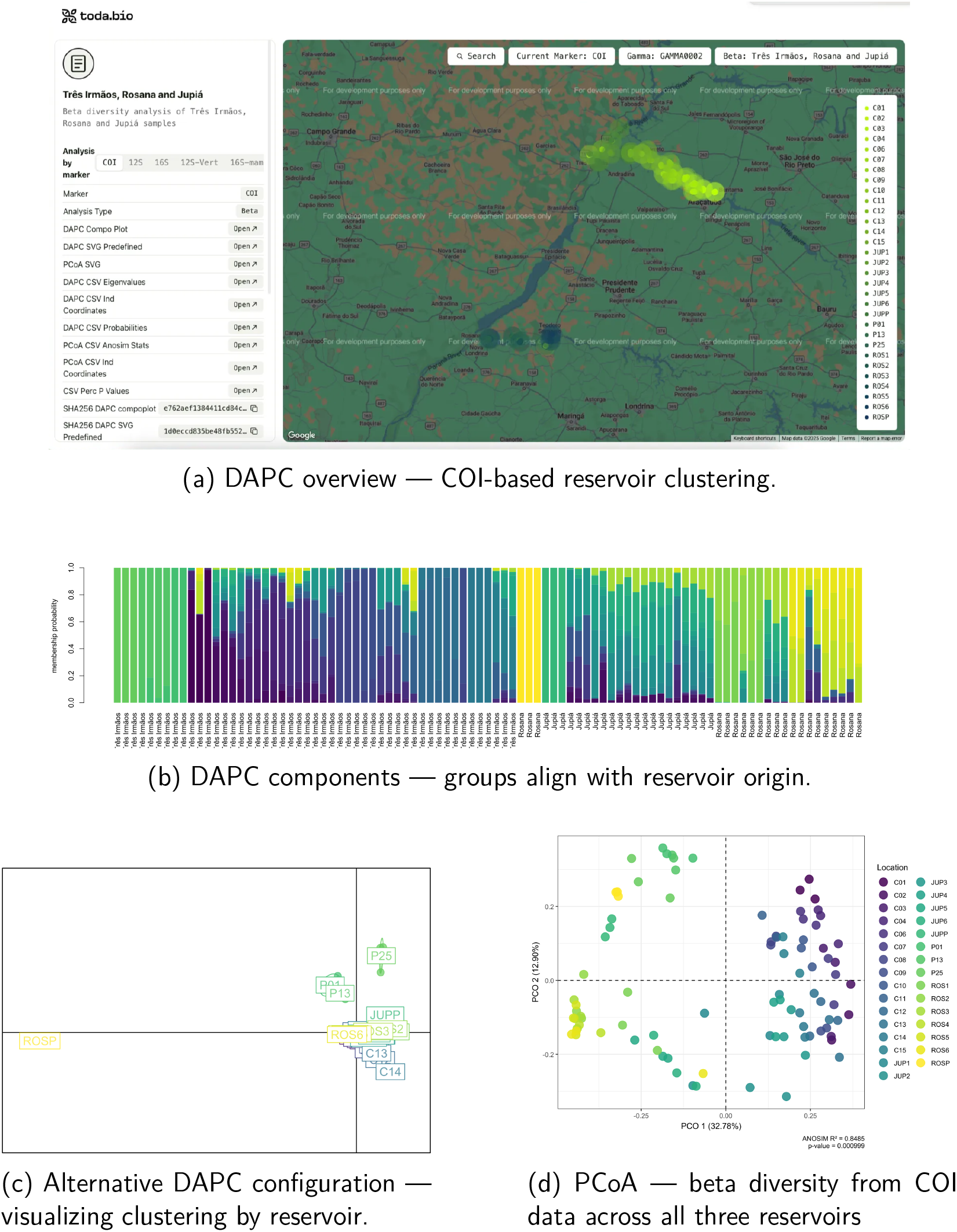
COI-based beta diversity analyses across 3 reservoirs. (a) Spatial DAPC overview of grouped samples; (b) DAPC component comparison shows consistent grouping by reservoir; (c) Alternative DAPC view; (d) PCoA supports clear reservoir separation. Samples from 31 stations (91 samples total) are grouped based on discriminant probability. Stations labeled C and P belong to Três Irmãos, JUP to Jupiá, and ROS to Rosana

**Figure 9.**
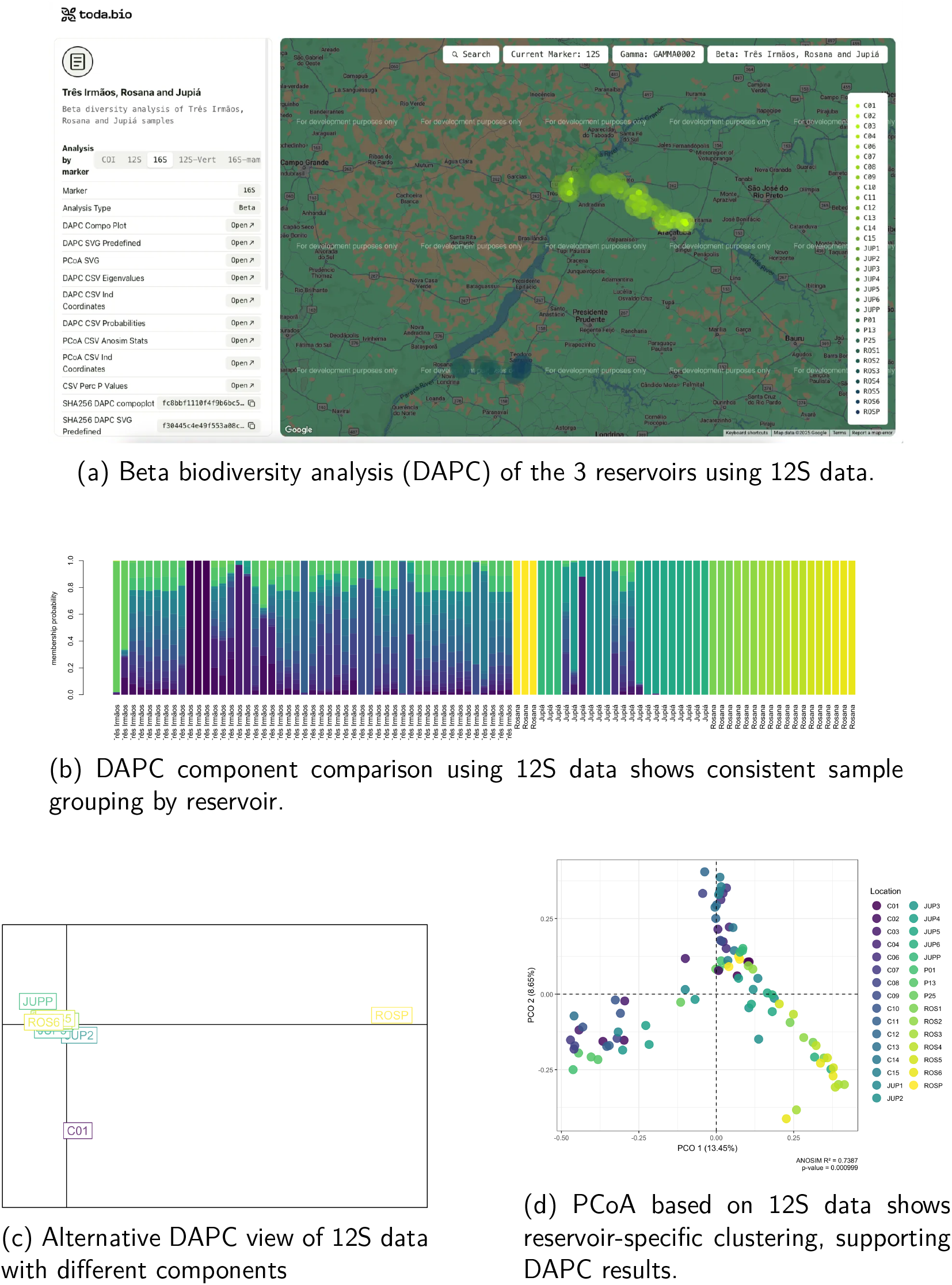
Beta diversity analysis of eDNA from three reservoirs using the 12S marker. DAPC and PCoA subfigures show consistent clustering by reservoir, revealing distinct biodiversity profiles. Samples (n = 91) from 31 stations are grouped by discriminant probability. Similar colors reflect reservoir-specific genetic signatures. Station codes: C and P = Três Irmãos, JUP = Jupiá, ROS = Rosana.

**Figure 10.**
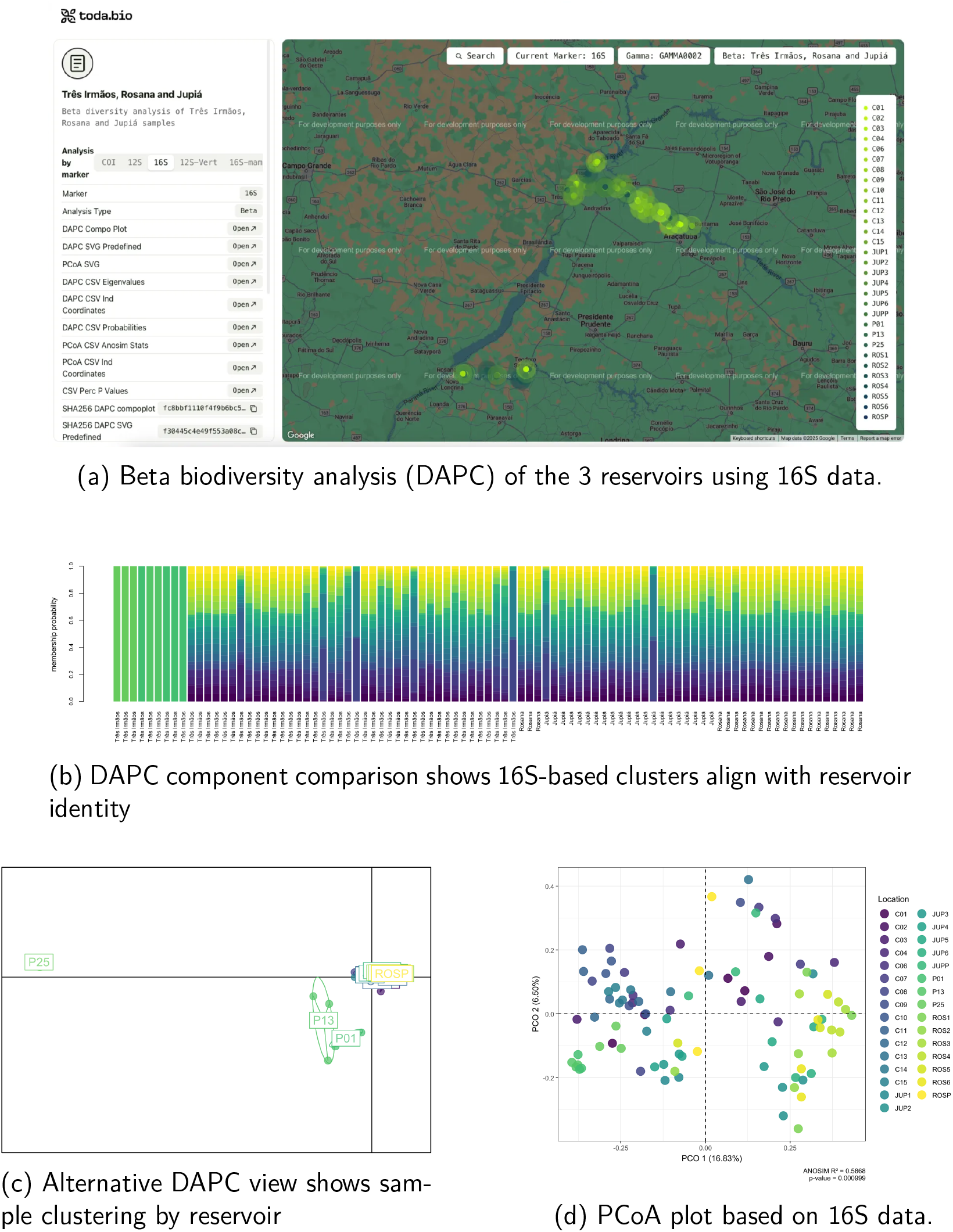
Beta diversity analysis of eDNA samples using the 16S marker shows consistent clustering by reservoir across 91 samples from 31 stations. Multivariate methods (DAPC and PCoA) reveal clear spatial separation, supporting the use of 16S for biogeographic attribution. Color-coded clusters correspond to reservoir identity: C and P = Três Irmãos; JUP = Jupiá; ROS = Rosana.

The supplementary material 6 is the machine-friendly biodiversity report in JSON format and supplementary materials 7 to 12 provide the full statistical out-puts used in the beta diversity analyses based on COI data. Each file name includes a SHA256 hash of its contents, ensuring integrity and enabling independent verification of the results.

Similar figures and outputs were generated for the 12S and 16S markers, but none showed the same degree of resolution and sample clustering as COI. For full statistical outputs used in the beta diversity analyses based on 12S and 16S data, please refer to supplementary materials 14-25.

Taken together, these results confirm that COI is the most effective marker for both biodiversity richness and reservoir differentiation, and is therefore the most suitable candidate for future applications of eDNA in proof-of-origin and biomonitoring frameworks. Among the three genetic markers used in this study (COI, 12S, and 16S), the cytochrome oxidase I (COI) gene proved to be the most informative for distinguishing biodiversity patterns between the Rosana and Jupiá reservoirs.

The COI marker consistently generated higher richness metrics across campaigns. In the third campaign (July 2024), it yielded 2,530 ASVs and 1,804 OTUs, compared to 449 ASVs / 357 OTUs for 12S and 778 ASVs / 416 OTUs for 16S (see Table 3). This higher resolution was evident in both taxonomic classification and the capacity to separate samples by reservoir of origin in multivariate analyses. The COI dataset also achieved broader species-level identification due to better representation in public reference databases such as BOLD and GenBank. For example, among classified ASVs in the third campaign, COI identified 83 fish species in Jupiá and 70 in Rosana, compared to lower counts in 12S and 16S (Supplementary Materials 5–7).

Moreover, ordination techniques such as Principal Coordinates Analysis (PCoA) and Discriminant Analysis of Principal Components (DAPC) revealed clearer clustering patterns when using only COI data. Samples grouped consistently by reservoir, indicating strong marker performance in capturing community structure and enabling proof-of-origin attribution.

While 12S and 16S remain valuable for complementarity—especially for taxa underrepresented in COI databases or for broad eukaryotic coverage—the superior resolution of COI supports its continued use as a primary marker in eDNA studies focused on freshwater fish biodiversity and sample provenance.

This result reinforces the practical benefits of COI for environmental monitoring programs, particularly in systems like the Paraná River Basin, where molecular tools are needed for regulatory compliance, conservation, and invasive species management.

## 3 Materials and Methods

### 3.1 Study Area

The study was conducted in two large reservoirs of the Paraná River Basin in southeastern Brazil: Jupiá and Rosana. Both are part of a cascade system of hydropower dams and are located in the transition between the Upper Paraná and Paranapanema sub-basins.

HPP Jupiá (officially Engenheiro Souza Dias) is a reservoir formed by damming the Paraná River downstream of Ilha Solteira. It has a surface area of approximately 330 km2 and receives inflows from major tributaries, including the Sucuriú River and the Três Irmãos Reservoir, in Tietê. It exhibits complex hydrodynamics due to floodplain interactions and a relatively large storage volume, which results in slow water renewal and extensive recirculation zones.

HPP Rosana, located downstream of HPP Taquaruçu on the Paranapanema River, is a smaller run-of-river reservoir with an area of approximately 220 km2. It has a more linear morphology and faster water turnover due to its operational characteristics. The reservoir’s hydrology is influenced by upstream dam releases and tributary inflows, but lacks significant lateral connectivity to floodplains.

Maps of sampling stations and hydrodynamic conditions for both reservoirs are provided in Figures 1a and 1b, and detailed sampling coordinates are available in Table 1.

### 3.2 Hydrodynamic Modeling

Hydrodynamic modeling was conducted to simulate flow velocity fields and water renewal dynamics in both reservoirs, allowing post hoc interpretation of eDNA transport and persistence.

For both Jupiá and Rosana, we applied a two-dimensional (2D) hydrodynamic model using a finite volume approach based on the Saint-Venant shallow water equations. Bathymetric data was provided by the concessionaire of the reservoirs and hydrometric data were sourced from the Brazilian National Electric System Operator (ONS) and local hydropower authorities. Simulations were conducted separately for the rainy and dry seasons of each sampling year.

Simulation outputs were used to generate velocity time series and instantaneous current fields for both reservoirs. These were integrated with spatial sampling data to evaluate whether hydrodynamic conditions influenced eDNA persistence and detectability, although no modeling information was used a priori for sampling design.

### 3.3 Sampling Design and eDNA Collection

After analyzing the data from both pilot samplings and the hydrodynamic modeling (see Results section), five new stations, named ROS2–ROS6 in Rosana and JUP2–JUP6 in Jupiá, were sampled in November 2023. Sampling stations were distributed along the entire reservoirs and covered each hydrodynamic compartment used to evaluate water renewal times across the Rosana and Jupiá rivers. In the third survey, the same six stations were sampled again in both reservoirs, and three additional stations were included in Rosana (ROSE1–ROSE3) to increase the representativeness of sampling during the dry season (July), when water velocity is high and can enhance eDNA transport and dilution. A total of 44 and 55 water samples were collected for Jupiá and Rosana, respectively, across the three surveys. At each station, water was sampled in three to five replicates, all collected 1 meter above the benthic bottom using a Van Dorn bottle. Replicates were collected within approximately 10-minute intervals. One liter of water was collected per replicate and stored in sterilized polypropylene bottles (Nalgon, Cat. No. 2330) at 8 °C until filtration through 0.22 μm Sterivex filters (Merck-Millipore, Cat. No. SVGP01050) in vaccuum pump. After filtration, residual liquid was expelled by injecting air, and the filters were stored at 4 °C in sterile Longmire’s buffer ([8] until DNA extraction. All non-disposable equipment was decontaminated with a 10% (v/v) sodium hypochlorite solution and rinsed with autoclaved distilled water to prevent cross-sample contamination.

### 3.4 DNA Extraction and Sequencing

DNA extraction from the Sterivex filters was performed using the DNeasy® Blood & Tissue kit (Qiagen) optimized for environmental samples with high organic load ([**?** ]). Filters were incubated with AL buffer, phosphate buffer, and proteinase K to maximize cell disruption and DNA yield. Extractions were carried out in a dedicated clean laboratory space to minimize contamination, with extraction blanks included in each batch. DNA quantification was conducted using a Qubit fluorometer with the dsDNA HS Assay Kit (Thermo Fisher Scientific). Extracted DNA was stored at *−*20°C until amplification and library preparation.

Amplification targeted three molecular markers: mitochondrial cytochrome oxidase I (COI), 12S ribosomal RNA, and 16S ribosomal RNA. The following primer sets were used:

- COI: Fishe-Mini-SH-E-F/Fishe-Mini-SH-E-R
- 12S: MiFish-U-F / MiFish-U-R
- 16S: 16S-uni-F / 16S-uni-R

Amplification was performed in 25 μl using Platinum™ Multiplex PCR Master Mix (Applied Biosystems). Thermocycling conditions was initial denaturation at 95 ° C for 2 minutes, 25 cycles at 95 ° C for 30 seconds, annealing at different temperatures for each primer pair for 90 seconds, and extension at 72 ° C for 30 seconds, followed by a final extension at 72°C for 10 minutes. Primers detailed in supplementary material 13.

For the library preparation, the amplicons were indexed using Nextera XT indexes (IDT for Illumina). Amplicons concentration was normalized to 4nM, amplicons were pooled in one library and purified with AMPure XP magnetic beads (Beckman Coulter). Library quality control was performed using Tapestation (Agilent). The sequence was performed on Illumina NovaSeq 6000 and NextSeq 1000 (2 *×* 150 bp paired-end). Sequencing runs included extraction control to monitor cross-contamination and barcode misassignment.

### 3.5 Bioinformatics and Statistical Analyses

After sequencing, raw sequence reads were evaluated using FASTQC [9] and MultiQC [9] . All generated fastq files were imported individually (forward and reverse) into the QIIME2 pipeline ([10]) for further quality filtering and taxonomic classification. Primer trimming, read denoising, merging the remaining paired reads into amplicons, and chimera removal steps were carried out with DADA2. The resulting Amplicon Sequence Variants (ASVs) were clustered into MOTUs (99% of identity among all samples ASVs sequences) with DADA2 ([11]) after the removal with blastn of sequences of contaminants from bacteria, vírus, humans, adapters, and primers, among others. The taxonomy identification was performed using BLASTn ([12]) searches in nt database (https://ftp.ncbi.nlm.nih.gov/blast/db/). 5 hits with an e-value lower than 1e-10 and a query cover higher than 95% were allowed. A series of scripts are executed to classify OTUs based on the top five BLASTn hits. Hits with identity and coverage greater than 98% and 95%, respectively, are considered for classification. The common ancestor (taxonomy retrieved from Environment for Tree Exploration (ETE) python toolkit ([13] among the classifications of the 5 hits is used to determine the final classification of the OTU in question. The final classification is then used to retrieve information via the GBIF API [14], including taxonomy, distribution, vernacular name, and Red List (IUCN) category ([7]). In the absence of high-confidence hits, assignments were made to the lowest available taxonomic level.

Heatmaps of the log10 frequencies of OTUs were constructed based on OTUs for fish species. OTUs frequencies were computed considering all three markers (CO1, 16S, and 12S) combined using lattice package R v. 3.3.2 ([15]).

Alpha diversity metrics (e.g., observed ASVs/OTUs, Shannon index (H’)) were calculated per sample using R ([16]). Beta diversity was assessed using Bray–Curtis dissimilarity and visualized through ordination methods including Principal Co-ordinates Analysis (PCoA) and Non-metric Multidimensional Scaling (NMDS) , implemented via the *vegan* package in R ([17]).

Discriminant Analysis of Principal Components (DAPC) was employed to assess whether samples could be correctly assigned to their source reservoir. Analysis of Similarities (ANOSIM) tested the statistical significance of clustering between reservoirs and campaigns.

Discriminant Analysis of Principal Components (DAPC) was employed to assess whether samples could be correctly assigned to their source reservoir, using the *adegenet* package in R ([18]). Analysis of Similarities (ANOSIM) tested the statistical significance of clustering between reservoirs and campaigns, as implemented in the vegan package [2].

Richness and taxonomic composition were compared across campaigns and between reservoirs using heatmaps, Venn diagrams, and stacked bar plots, all generated in R using packages including *phyloseq* ([19], *vegan* ([17], and *ggplot2* ([20]. Supplementary Materials 5–9 include detailed classifications per sample and marker.

### 3.6 Machine-friendly automated reports with verifiable integrity and proof of origin

All sequencing data, bioinformatics intermediate files and final outputs, were organized into machine-readable biodiversity reports to ensure reproducibility, transparency, and independent verification.

The reporting pipeline begins with raw sequence data (FASTQ files), which are analyzed through a standardized workflow using Nextflow, Python, and R scripts.

Each script generates a specific output (e.g., filtered reads, ASVs, OTUs, taxonomic tables), and all input–output relationships are recorded. The final outputs are consolidated into a single JSON file that includes:

- Sample metadata (timestamp, location, sample type)
- URLs of input and output files (stored on AWS and IPFS)
- Logs of analytical steps and parameters
- Alpha and beta diversity metrics
- Hashes (SHA256) of all files and the report itself

This JSON file functions as a **machine-friendly biodiversity report** — readable by both humans and systems — and is permanently stored on the InterPlanetary File System (IPFS). Its IPFS address (a hash of the content) guarantees immutability.

The hash of the JSON report is recorded on the Bitcoin-based Liquid Network blockchain as a digital asset with the following properties:

- Non-fungible (precision = 0, amount = 1)
- Not reissuable (ensuring uniqueness)
- Linked to a specific report and sampling event

The resulting asset serves as a verifiable, tamper-proof record of the biodiversity analysis, enabling the traceability of the sample’s origin and analytical history without reliance on trust in any institution.

This system provides an auditable chain of custody for biodiversity data, fulfilling regulatory demands while enabling interoperability across platforms. Clients may integrate reports via JSON APIs, visualize them on dashboards, or tokenize them on secondary blockchains for use in biodiversity credit markets.

This approach transforms biodiversity reporting from a static PDF into a transparent, traceable, and trustless verification mechanism.

## 4 Conclusion

This study demonstrates that environmental DNA (eDNA) profiles can effectively distinguish the geographic origin of water samples collected from large aquatic ecosystems. By integrating standardized multi-marker metabarcoding with supervised classification and decentralized data storage, we provide a framework for generating verifiable biodiversity reports that are permanently registered and auditable via blockchain infrastructure. This approach introduces a transparent, scalable, and trustless method for environmental traceability. Beyond expanding the applications of eDNA, our findings offer a novel foundation for independently verifying sample provenance in biodiversity monitoring and ecological compliance workflows.

## Acknowledgments

We thank Angelica Beccato, Gabriela Monteiro, Arthur Rotiroti, Cristiano Laluce, and André Silva for their invaluable contributions. Their insights into the sampling area and prior experience with fisheries management greatly enriched this work. We are also grateful for their careful reading of the manuscript, which helped improve its clarity and rigor. We are also thankful to Maya Parbhoe, Adolfo Contreras and Valerio Vaccaro for their assistance with token issuance.

## Data Availability Statement

All raw sequencing data, processed outputs (ASVs, OTUs, taxonomic classifications), and biodiversity metrics are available as machine-readable JSON reports stored on IPFS. Each report includes metadata, analysis logs, and SHA256 hashes, with provenance registered on the Liquid Network blockchain.

## 6 Supplementary Materials

**SUP01** – [12S mifish classification 3rd campaign S5](https://drive.google.com/file/d/1lOfsLBug2kDlglWHWDCIE9R-WHEE_ubP/view?usp=drive_link)

**SUP02** – [16S UNI classification 3rd campaign S6](https://drive.google.com/file/d/1iM6m2dcAJu3so76wWpF32fxFCWR2W0ZO/view?usp=drive_link)

**SUP03** – [COI classification 3rd campaign S7](https://drive.google.com/file/d/1L2e8MmWjWSiVJNmMbYHSv7UudIRZts87/view?usp=drive_link)

**SUP04** – [Jupiá community combinations](https://drive.google.com/file/d/1eNMcYy2z0AlHSvKZjj6kM3x_fFYSV2Wf/view?usp=sharing)

**SUP05** – [Rosana community combinations](https://drive.google.com/file/d/1P2ghJeNi8vV1FruH7vHsx4fN0cHw1tpE/view?usp=sharing)

**SUP06** – [Beta diversity COI (JSON report)](https://drive.google.com/file/d/1sL7FdyIDWrAI5MAuiqptZA3kYTDg_RdR/view?usp=sharing)

**SUP07** – [COI DAPC probabilities](https://drive.google.com/file/d/1B59hyaKzXaXNaPQzQoDtcluEFDd0Rpft/view?usp=sharing)

**SUP08** – [COI DAPC individual coordinates](https://drive.google.com/file/d/1cl6R1R95PA1bCPvVLKdTkzHr4OIT_j-h/view?usp=sharing)

**SUP09** – [COI DAPC eigenvalues](https://drive.google.com/file/d/1G8GLVRPuBQ0gQ8G-kNP29VYGj_pqQHIS/view?usp=sharing)

**SUP10** – [COI PCoA and ANOSIM statistics](https://drive.google.com/file/d/1beizJRIUaOTvCM6UAZFd9OeW-VikrbXp/view?usp=sharing)

**SUP11** – [COI PCoA individual coordinates](https://drive.google.com/file/d/14ShyKyEUQd2jc7u-srmQjPls3ZjJ9MHR/view?usp=sharing)

**SUP12** – [COI permutation p-values (0.05)](https://drive.google.com/file/d/13PxwTsolgjmY0egBFQ_u21MWvE5a8IiJ/view?usp=sharing)

**SUP13** – [Primer sequences used](https://drive.google.com/file/d/13P3HMdWx2qOzeHdwHIBd_9iMAo1nj2em/view?usp=sharing)

**SUP14** – [12S DAPC probabilities](https://drive.google.com/file/d/1O07GMlaSz0LXQ_g203JIxn4lW2oN1eu9/view?usp=sharing)

**SUP15** – [12S DAPC individual coordinates](https://drive.google.com/file/d/1hBA0LToWlNln2cMECbG1cErCmFg_n2dc/view?usp=sharing)

**SUP16** – [12S DAPC eigenvalues](https://drive.google.com/file/d/1qJYFC48oIE519TnJ-aGHwFfTw57Owing/view?usp=sharing)

**SUP17** – [12S PCoA and ANOSIM statistics](https://drive.google.com/file/d/1iyuKMe0nkbZgLm3lzNPQKWNk4mV3GJue/view?usp=sharing)

**SUP18** – [12S PCoA individual coordinates](https://drive.google.com/file/d/1pfuv-wpTAG60beoZwfkhHnEPq--Y0MJd/view?usp=sharing)

**SUP19** [– 12S permutation p-values (0.05)](https://drive.google.com/file/d/156Y4eAKBw3oEv_4sX_McuMmMEb3o75YP/view?usp=sharing)

**SUP20** – [16S DAPC probabilities](https://drive.google.com/file/d/1hqrnTcyjj_dyjy62WRZ2BZcEkZ1d7F6Q/view?usp=sharing)

**SUP21** – [16S DAPC individual coordinates](https://drive.google.com/file/d/1YooWLHQ-IKfAmoUMBS9X38VZ6uzXKcqe/view?usp=sharing)

**SUP22** – [16S DAPC eigenvalues](https://drive.google.com/file/d/1u4EVn0BE-Gbm2pakATc7-gJ0Iwbw5yZJ/view?usp=sharing)

**SUP23** – [16S PCoA and ANOSIM statistics](https://drive.google.com/file/d/1dCHBko5O42_oD6f7UcOIGf_aL0iW12SE/view?usp=sharing)

**SUP24** – [16S PCoA individual coordinates](https://drive.google.com/file/d/1sraWth8bQrsAXCVPnXV-sLeSZguGk9nY/view?usp=drive_link)

**SUP25** – [16S permutation p-values (0.05)](https://drive.google.com/file/d/1kODBUwlU1qN_SRfxMIyY2fj21H5P-Hmc/view?usp=sharing)

